# Parsimonious identification of evolutionary shifts for quantitative characters

**DOI:** 10.1101/184663

**Authors:** Olivier Chabrol, Gilles Didier

## Abstract

Detecting shifts in the rate or the trend of a character evolution is an important question which has been widely addressed. To our knowledge, all the approaches developed so far for detecting such shifts from a quantitative character strongly involved stochastic models of evolution.

We propose here a novel method based on an asymmetric version of the linear parsimony (aka Wagner parsimony) for identifying the most parsimonious split of a tree into two parts between which the evolution of the character is allowed to differ. To this end, we evaluate the cost of splitting a phylogenetic tree at a given node as the integral, over all pairs of asymmetry parameters, of the most parsimonious cost which can be achieved by using the first parameter on the subtree pending from this node and the second parameter elsewhere. By testing all the nodes, we then get the most parsimonious split of a tree with regard to the character values at its tips.

A study of the partial costs of the tree enabled us to develop a polynomial algorithm for determining the most parsimonious splits and their evolutionary costs. Applying this algorithm on two toy examples shows that using more than one asymmetry parameter does not always lower parsimonious costs. By using the approach on biological datasets, we obtained splits consistent with those identified by previous stochastic approaches.

## 1 Introduction

Changes in the rate and/or the trend of a character evolution are often associated with major biological phenomenons such as adaptation or evolutionary convergence [10]. Detecting shifts in the evolution of a trait is thus an important question, which has been widely addressed, notably in the case of quantitative characters. All the approaches developed so far for studying this question are based on stochastic models of evolution for continuous characters, mainly Brownian motion or Ornstein-Uhlenbeck processes [1, 9, 3, 6, 11, 10, 13, 14, 15, 16, 17]. The method presented in [2] used parsimony approaches but only for reconstructing ancestral states which were next compared with simulations under Brownian models. Other approaches addressed the related question of testing the significance of splits given *a priori* (e.g. from environmental or dietary considerations) with regard to stochastic models [21, 22, 18].

We propose here a novel approach based on an asymmetric version of the linear parsimony introduced in [4] and further studied in [5], in both cases in an ancestral reconstruction context. We do not discuss the pros and cons of parsimonious approaches with regard to those based on stochastic models [19, 12, 8]. From a technical point of view, ideas behind parsimonious algorithms can generally be transposed to stochastic frameworks. We plan to adapt principles used in our computations for determining the most likely split of a phylogenetic tree with regard to Brownian and Ornstein-Uhlenbeck models.

The problem that we are studying here can be formally stated as follows. Being given a phylogenetic tree and a continuous character which is known only for extant species (i.e. here the tips of the tree), we aim to identify the node of the tree from which the evolution of the character differs (or differs the most) from the rest of tree, if such a node exists.

A parsimonious framework enables us to compute the minimal evolutionary cost of the extant character values. This cost plays the role of the likelihood in probabilistic models. In the asymmetric linear parsimony case, the evolutionary cost depends on the asymmetry parameter *α*. In order to evaluate the relevance of putting an evolutionary shift at a node *n*, we consider evolutionary costs obtained by using an asymmetry parameter on the subtree 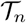 and another parameter on the rest of the tree. Testing all the nodes of the tree enables us to identify the subtree on which the evolution of the character differs the most. We first provide a polynomial algorithm for computing such costs and for integrating their over all pairs of asymmetry parameters. This way we get a parsimonious method for finding the subtree on which the evolution differs the most. We show on toy examples that considering more than one asymmetric does not always lead to lower the evolutionary cost. Finally, our approach is applied on two previous biological datasets from [18] and [20]. Shifts detected with our approach are consistent with those obtained from methods based on stochastic models and well separate clades with specific evolution like whales from other cetaceans.

We developed *ParSplit* a software performing the parsimonious split of a phylogenetic tree with regard to extant values of characters written in C language. Sources of the software and toy examples are available at https://github.com/gilles-didier/ParSplit.

The rest of the paper is organized as follows. Section 2 presents definitions, notations and asymmetric linear parsimony then outlines the approach. The *partial cost functions*, which are the bases of our computations, are defined and studied in Section 3. Next, Section 4 shows how to integrate the (split) costs over all the possible values of the asymmetry parameter(s). Finally, results obtained with our approach on two toy examples and on two biological datasets are presented in Section 5 which ends with a short discussion.

## 2 Asymmetric parsimony

### 2.1 Definitions and Notations

We use the same notations as [5]. The cardinal of a finite set 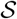 is noted 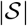. In what follows, 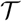 designates a rooted tree which may or may not be binary. As it should lead to no confusion, we still write 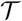 for its set of nodes. For all nodes 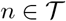, we put

- 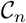 for the set of child nodes of *n*,
- *τ_n_* for the length of the branch ending at *n*,
- 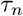 for the subtree of 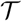 rooted at *n*.

Let us consider a subset 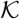 of nodes of 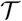 and a map *ϑ* from 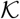 to the set of real numbers 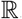. The map *ϑ* will be referred to as the *initial function* and the nodes of 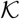 are said *known*. For all nodes *n* of 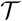, we put 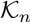 for the subset of known nodes of the subtree 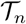, i.e. 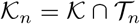. Though the general case can be treated as in [5], we assume that all and only the tips of the tree have known states in order to lighten the statements. In plain English, 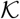 (resp. 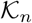) is the set of tips of 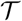 (resp. of 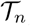).

The values of 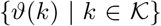 are the *known states* of 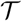. For all nodes *n*, we put 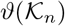 for the set 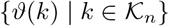.

A *ϑ-assignment* of 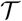 is a map *ξ* from 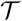 to the set of real numbers which extends *ϑ* (i.e. such that 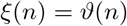 for all nodes 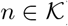. The set of all *ϑ*-assignments of 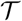 is noted 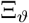.

### 2.2 Asymmetric parsimonious cost

A parsimony framework is based on a way of computing the evolutionary cost of an assignment. In the time-dependent-asymmetric-linear parsimony (*TDALP*, see [5]) case, an ancestor/child transition from value *x* at node *n* to value *y* at its child *m* is associated with the cost:

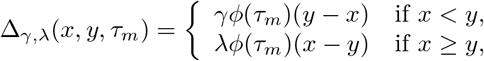

where *λ* and *γ* are two nonnegative real numbers and *ϕ* is a function from 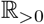 to 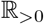 (in what follows, we make the assumption that *τ_n_* > 0 for all nodes 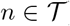). The case where *ϕ* is constant with *ϕ(τ)* = 1 for all *τ* corresponds to the standard parsimony scheme [4, 7]. In an evolutionary context, it makes sense to choose a decreasing function *ϕ*, but any positive function can be used. In the current implementation of the approach, users can choose between 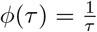 and *ϕ(τ)* = 1.

The cost Δ*_γ,λ_*(*ξ*) of an assignment *ξ* of 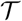 is then the sum of all the costs of its ancestor/child transitions:

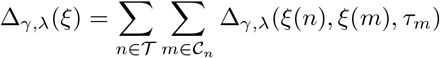

Let us first remark that multiplying both *λ* and *α* by a positive constant factor just lead to multiply the assignments’ cost by the same factor. Since computing an evolutionary cost with *α* = *λ* = 0 makes no sense, we can assume *γ* + *λ* > 0 and divide both parameters by *γ + λ* in order to normalize the parsimonious costs. From now on, we assume that *γ* + *λ* = 1. The cost of an assignment then only depends on a single parameter *α* ∈ [0, 1], and is called *α-cost*:

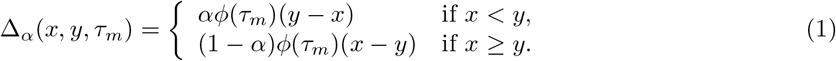

Below, *α* will be referred to as the *asymmetry parameter*. Reconstructing with 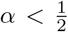 (resp. with 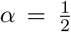, with 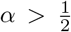) makes the cost of an increase smaller than (resp. equal as, greater than) that of a decrease of the same amount. Intuitively, it corresponds to the assumption that the character evolves with a positive trend (resp. without trend, with a negative trend). Note that a slightly different parametrization is considered in [5].

The *(parsimonious) α-cost* of the pair 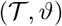 is defined as the minimal cost which can be achieved by a *ϑ*-assignment:

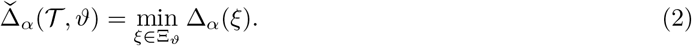

### 2.3 Outline of the approach

Our aim is to find the most parsimonious way of splitting a phylogenetic tree 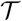 into two parts according to an initial function *ϑ* (i.e., the characters values of tips). In what follows, we will consider the two ways of splitting a phylogenetic tree at a node *n* displayed in Figure 1 and below referred to as *A-split* and *B-split*. Devising a parsimonious approach for identifying evolutionary shifts first requires to be able to associate an evolutionary cost at a given split.

**Figure 1:**
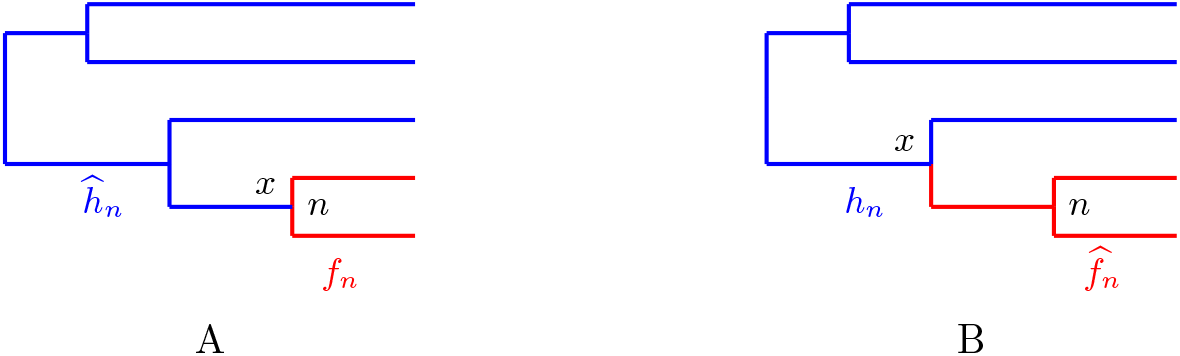
Two different ways of splitting a phylogenetic tree at a node *n*.

In a parsimonious context, a natural idea should be to define the cost of a split of a tree into two parts as the minimum cost, over all pairs (*α, α'*) ∈ [0, 1]^2^, which can be achieved by using parameter *α* on a part of the tree and parameter *α'* on the other part. Unfortunately, even without splitting the tree, minimizing the parsimonious cost over all parameters *α* leads to trivial solutions: by setting *α* = 0 (resp. *α* = 1) the assignment which associates the greatest (resp. the smallest) of the known values to all unknown nodes has a null *α*-cost!

In order to overcome this issue, we first define the *no-split cost* of 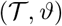 as the sum over all parameters *α* ∈ [0, 1] of the *α*-parsimonious costs 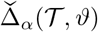. Note that this cost is parameter-free and depends only on 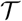 and *ϑ*. It somehow reflects the disparity of the character with regard to the tree. We then define the cost of a split of 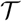 into two parts as the sum over all pairs of parameters (*α, α'*) ∈ [0, 1]^2^ of the smallest cost which can be achieved by using parameter *α* on the first part and parameter *α’* on the second one. The most parsimonious split of a phylogenetic tree with regard to a given initial function is either the whole tree if the no-split cost is lower than all the costs of the *A*- and *B*-splits, or the *A*- or *B*- split with the smallest cost otherwise.

The next two sections show how to compute the no-, A- and B-splits costs.

## 3 Partial cost functions

For all nodes *n* of 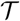, we shall define and study the partial cost functions 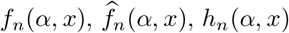 and 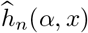 that give the minimal costs which be obtained by associating *x* to *n* or its direct ancestor and by using parameter *α* on the corresponding parts of the tree as displayed in Figure 1.

Namely, following [5, 4], for all nodes *n* of 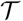, we put *f_n_*(*α, x*) for the minimal cost which can be achieved by an assignment *ξ* of the subtree 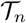 (red part of Figure 1-left) such that *ξ*(*n*) = *x*. For all nodes *n* of 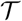 but its root, we put 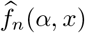 for the minimal cost which can be achieved by an assignment *ξ* of the tree consisting of *m*, the direct ancestor of *n* and of the subtree 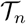, which will be referred to as the *stem subtree of n* and noted 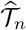 (red part of Figure 1-right), verifying *ξ*(*m*) = *x*.

Cost functions *h_n_*(*α, x*) and 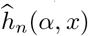 give the complementary costs of 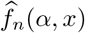 and *f_n_*(*α, x*). Namely, for all nodes *n* of 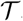, we put *h_n_*(*α, x*) for the minimum reconstruction cost of 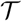 without *n*, the branch bearing *n* and the subtree rooted at *n* (blue part of Figure 1-right), which associates *x* to the direct ancestor of *n*. Last, we put 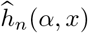 for the minimal cost which can be achieved by an assignment *ξ* of the whole tree with *n* and the branch bearing *n* but without the subtree rooted at *n* (blue part of Figure 1-left) which is such that *ξ*(*n*) = *x*.

We first recall and slightly adapt a Theorem from [5].

### Theorem 1 ([5]).

*Let n be an unknown node of* 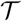. *The maps f_n_ and* 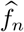 *are piecewise-linear and continuous and, for all α >* 0*, the maps x → f_n_*(*α, x*) *and* 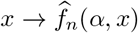 *are both convex*.

*More precisely, if all the nodes of* 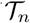 *are unknown then f_n_*(*α, x*) = 0 *for all α* > 0 *and all x. Otherwise, there exist:*

- *an integer u_n_ and a strictly increasing positive real sequence* 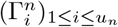,
- *an integer sequence* 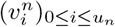 *and for, all* 0 *≤ i ≤ u_n_, a sequence* 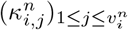 *of known states of* 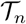 *(i.e. in* 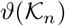*) verifying* 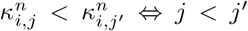; *by convention we set* 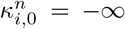 *and* 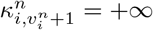,
- *two nonnegative real bi-sequences* 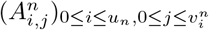 *and* 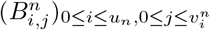,
- *two real bi-sequences* 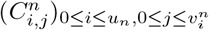 *and* 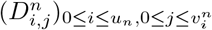
 *such that, by setting* 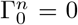 *and* 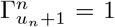 *and for all* 0 *≤ i ≤ u_n_, all* 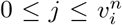, *all* 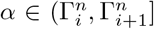 *and all* 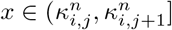, *we have*

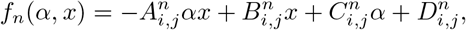

*all the coefficients being such that f_n_ is continuous. Moreover, the sequence* 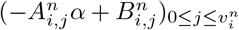 *(i.e. the x-coefficients of f_n_) is increasing with* 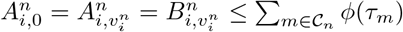 *and* 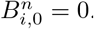.

*In the same way, if all the nodes of* 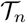 *are unknown then* 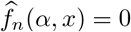 *for all α >* 0 *and all x. Other wise there exist* 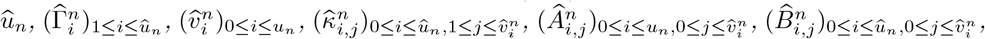 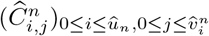, 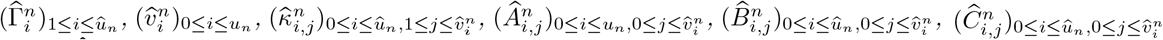 *and* 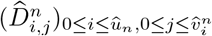 *verifying the same properties as their f_n_-counterparts except that we have* 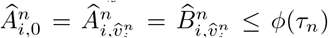 *and* 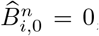 *and such that, for all* 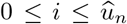, *all* 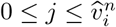, *all* 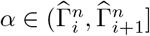 *and all* 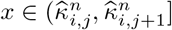, *we have*

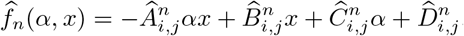

#### Proof

In [5], this theorem was established by considering another parametrization of the parsimony cost, namely

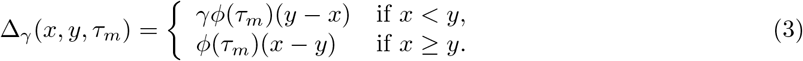

The version above was obtained by dividing the cost just above and the corresponding partial cost functions by *γ* + 1 and by applying the substitution 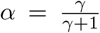 to transform the coefficients and the bounds of these cost functions.

A similar result can be stated for *h_n_* and 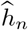.

### Theorem 2

*Let n be a non-root unknown node of* 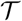. *The maps h_n_ and* 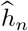 *are piecewise-linear and continuous and, for all α* > 0*, the maps x → h_n_*(*α, x*) *and* 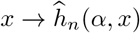 *are both convex*.

*More precisely, if all the nodes of* 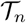 *are unknown then h_n_*(*α, x*) = 0 *for all α* > 0 *and all x. Otherwise, there exist:*

- *an integer *w_n_* and a strictly increasing positive real sequence* 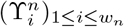,
- *an integer sequence* 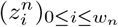 *and for, all* 0 *≤ i ≤ w_n_, a sequence* 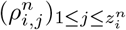 *of known states of* 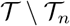 *(i.e. in* 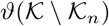*) verifying* 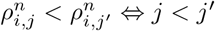; *by convention we set* 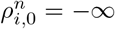 *and* 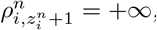,
- *four real bi-sequences* 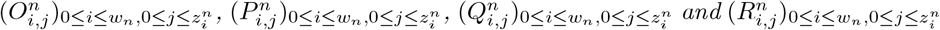 *such that, by setting* 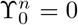 *and* 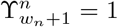 *and for all* 0 *≤ i ≤ w_n_, all* 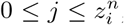, *all* 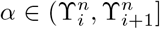 *and all* 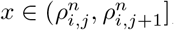, *we have*

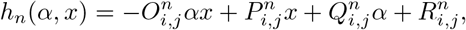

*all the coefficients being such that h_n_ is continuous. Moreover, the sequence* 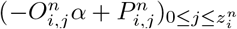 *(i.e. the x-coefficients of h_n_) is increasing with* 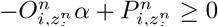 *for all* 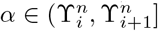

*In the same way, if all the nodes of* 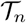 *are unknown then* 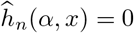 *for all α >* 0 *and all x. Other-wise there exist* 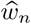, 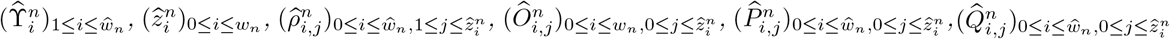 *and* 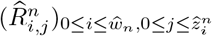 *verifying the same properties as their h_n_-counterparts and such that, for all* 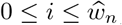, *all* 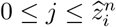, *all* 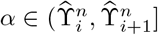 *and all x* 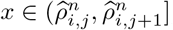, *we have*

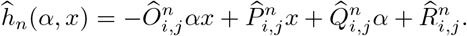

### Proof

The proof follows the same ideas as that of Theorem 1 from [5], a noticeable difference being that here the induction goes from the root to the tips. Namely, we shall prove the theorem by proving the three following assertions:

1. if *n* is a child of the root then *h_n_* satisfies the properties of the theorem;
2. for all nodes *n*, if *h_n_* satisfies the properties of the theorem then so it is for 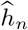;
3. for all nodes *n*, if 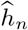 satisfies the properties of the theorem then so it is for all *h_k_* with *k* child of *n*.

By construction, if *n* is a direct descendant of the root *r*, we have that

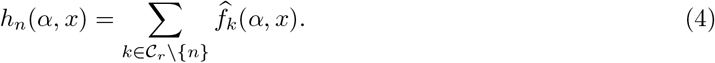

Assertion 1 then follows from Theorem 1.

In order to prove Assertion 2, let us remark that for all nodes *n*, by setting

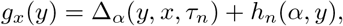

we have

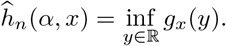

Let us assume that *h_n_* satisfies the properties of the theorem. In particular *h_n_* is piecewise linear with, for all 0 *≤ i ≤ w_n_*, all 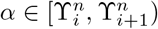, all 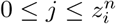 and all 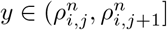:

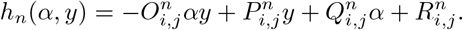

Let us first consider the case where *y ≥ x* for which we have Δ*_α_*(*x, y, τ_n_*) = (1 − *α*)*ϕ*(*τ_n_*)(*y − x*).

For all 0 *≤ i ≤ w_n_*, all 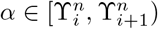, all 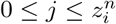 and all 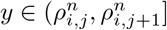, we get that

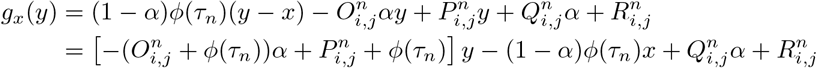

Since *h_n_* satisfies the properties of the theorem, it is convex with respect to *x*. The sequence 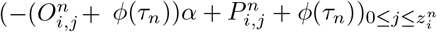 thus increases with *j*. Let 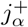 be defined as

- 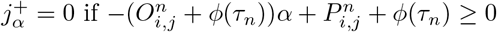 for all 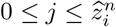;
- the greatest integer smaller or equal to 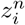 and such that 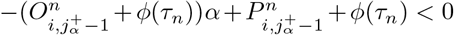 otherwise. In other words, 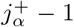 is the index of the last interval on which *g_x_* strictly decreases with *y ≥ x* and is the greatest index such that 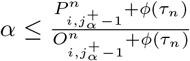.

Since *g_x_* decreases before 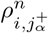 and increases after this value, we get that

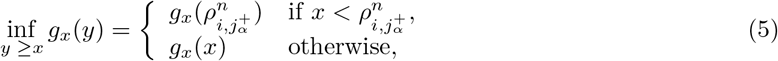

and that

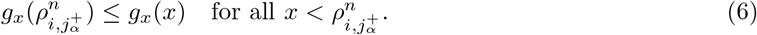

Let us now consider the case where *y < x* for which we have 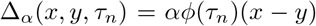. For all 0 ≤ *i < w_n_*, all 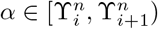, all 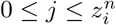 and all 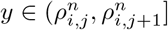, we get that

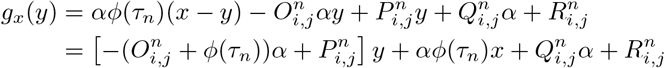

Let us define 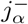 as

- 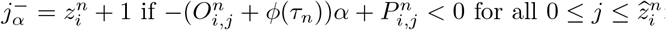;
- the smallest index such that 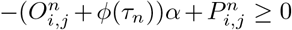 otherwise. In other words, 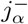 corresponds to the first interval on which *g_x_* does not decrease with *y < x* and is the smallest index such that 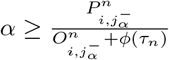.

We have that

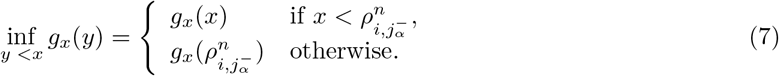

and that

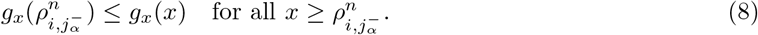

Since *ϕ*(*τ_n_*) is positive, 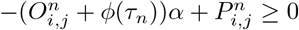 implies that 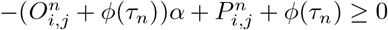 for all *i* and *j*. It follows that that

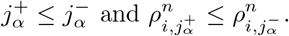

The fact that

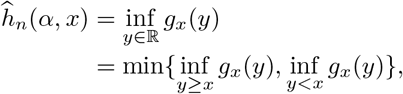

Equations 5 and 7 and Inequalities 6 and 8 imply that

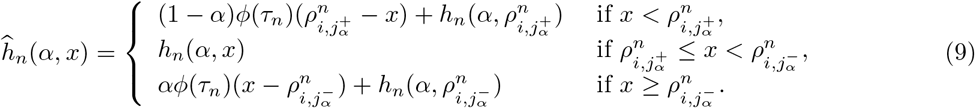

We assume here that *h_n_* satisfies the properties of the theorem. This first implies that for all *α*, the map 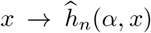 is piecewise linear and continuous. Next, the map *x → h_n_*(*α, x*) is convex, in particular between 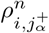 and 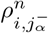. Moreover, definitions of 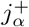 and 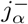 ensure that if 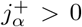 then 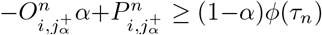 and if 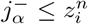 then 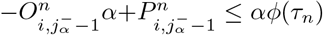. It follows that the map 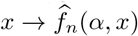 is still convex. Moreover, if 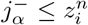 then the *x*-coefficient of 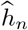 for 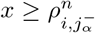 is *αϕ*(*τ_n_*) which is nonnegative. Otherwise, we have in particular that 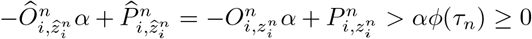, which implies that the *x*-coefficient of 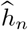 for 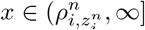 is positive.

Let us put Φ_1_, …, Φ*_p_* for the elements of

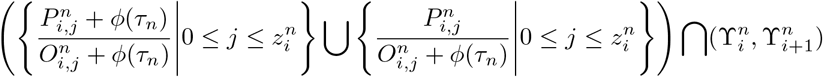

indexed in increasing order. By construction, the indexes 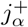 and 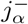 are both constant over all the sub-intervals 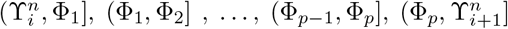. The fact that *h_n_* satisfies the properties of the theorem and Equation 9 imply that for all values *x*, the map 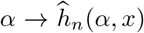 is linear over all the sub-intervals above. We get that the map 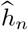 is piecewise linear and its continuity with regard to *α* is straightforward to verify at all bounds Φ*_k_* for 1 ≤ *k* ≤ *p*.

Last, Assertion 3 follows straightforwardly from Theorem 1 and from the fact that, for all children *k* of *n*, we have

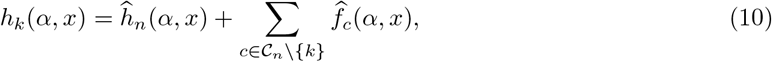

which ends the proof.

Theorem 2 suggests the procedure sketched in Algorithm 1 for computing the functions *h_n_* and 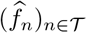 for all non-root nodes *n*.

Like the algorithm computing (*fn*)*_n∈T_* and 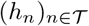 provided in [5], the complexity of Algorithm 1 depends on the total number of bounds (over asymmetry parameters and over character values) required for defining (*h_n_*)*_n∈T_* and 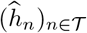. The number of bounds over character values, below referred to as *x-bounds*, is, by construction, smaller than 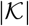 (we don’t count the two extrema −∞ and +∞). We shall bound the number of intervals over asymmetry parameters, below referred to as *α-bounds*, following the same ideas as in [5].

From now on, we make the implicit assumption that all bounds of the partial cost functions are actually required to define it. Namely, under the notations of Theorem 2 and for all nodes *n* of 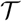, the cost function *h_n_* is such that:

- for all 0 ≤ *i* ≤ *w_n_* and all 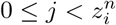, we have that

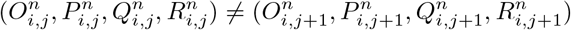
- for all 0 ≤ *i* < *w_n_*, there exists 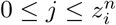 and 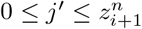 such that

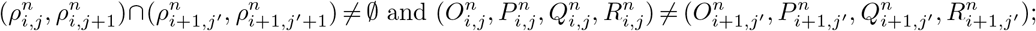
 and that the same holds for the partial cost functions 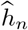, *f_n_* and 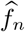.

Remark that since *h_n_* is continuous, the first assertions implies that for all 0 ≤ *i* ≤ *w_n_* and all 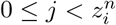, we have that

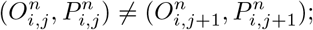

### Lemma 1

*Let n be an unknown node of* 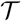. *For all* 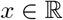, *the x-coefficient of h_n_ (resp. of* 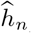*) decreases with γ*.

### Proof

We shall follow the same three steps as for the proof of Theorem 2 and the same ideas as for that of Lemma 1 from [5]. Let us first recall that Lemma 1 from [5] ensures that the *x*-coefficient of 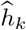 decreases with *α* for all children *k* of the root different from *n*. Actually, a slightly different parametrization is used in [5] but the same change of variable as for Theorem 1 shows that this lemma remains true with the parametrization used here.

First, for all children *n* of the root of 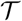, Equation 4 and Lemma 1 from [5] ensure that the lemma holds for *h_n_*.

Second, let us prove that if the lemma is true for *h_n_* then it is true for 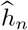. Let *α* ∈ [0, 1] be an asymmetry parameter and *i* be the index such that 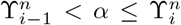. From the proof of Theorem 2, there exists an index 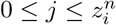 such that, by setting 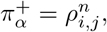, the *x*-coefficient of 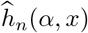 is equal to 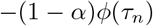 for 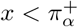 and greater than −(1 − *α*)*ϕ*(*τ_n_*) otherwise (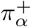 is possibly equal to −∞).

**Algorithm 1:**
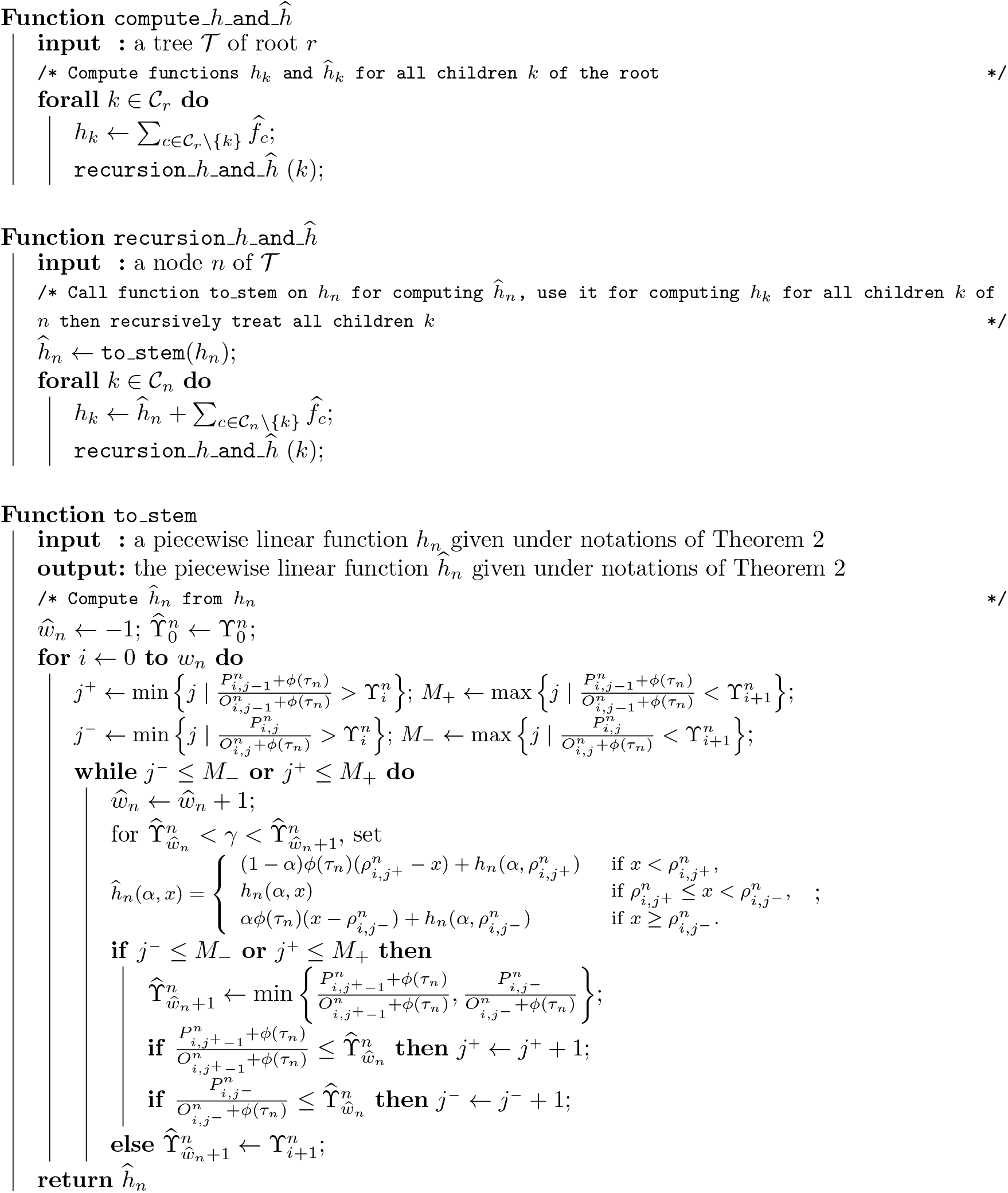
Computation of *h_n_* and 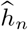 for all non-root nodes *n* of 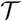.

Symmetrically, there exists an index 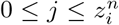 such that, by setting 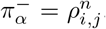, the *x*-coefficient of 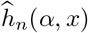 is equal to *αϕ*(*τ_n_*) for 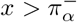 and smaller than *αϕ*(*τ_n_*) otherwise.

By construction and from the induction hypothesis, we get that both 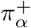 and 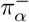 increase with *α*.

Let us now consider two asymmetry parameters 0 ≤ *α'* ≤ *α''* ≤ 1. We then have 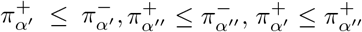, and 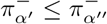. The first two inequalities come from construction (see the proof of Theorem 2) and the two last ones from the remark just above. This leaves only two cases to investigate:

1. 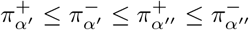 and
2. 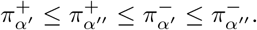.

By checking all the possible positions of *x* with regard to the bounds 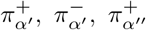 and 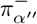 like in the proof of Lemma 1 of [5], we verify that the *x*-coefficient of 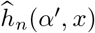 is always greater than that of 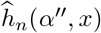 either directly or from the induction hypothesis both in Cases 1 and 2. The *x-coefficient* of 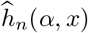 does decrease with *α*.

Third, we remark that if the *x-coefficient* of 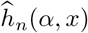 decreases with a then, since the same holds for all children *k* of *n* from Lemma 1 of [5], Equation 10 implies that the x-coefficient of 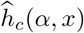 decreases with a for all children *c of n*.

### Proposition 1.

*Under the notations of Theorem 2 and for all nodes n of* 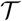, *we have that*

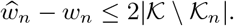

### Proof

Let *n* be a node of 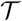 with 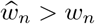. There exists an index *l* in 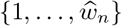 such that 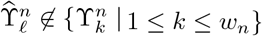 *1 ≤ k ≤ w_n_}*. Let *i* be the index such that 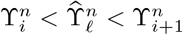. From the proof of Theorem 2, there exists an index 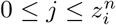 such that at least one of the following assertions holds:

1. 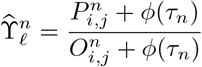
2. 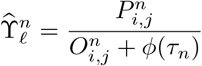

In other words, there are two types of *α-bounds* of 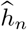 which are not *α-bounds* of *h_n_*.

If 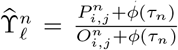 (Type 1) and still from the proof of Theorem 2, we have that 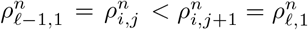. Lemma 1 then ensures that for all i’ > i and all j’ such that 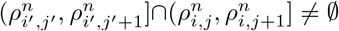, we have that 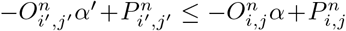 for all 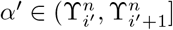 and all 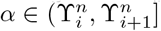 with α’ > α. It follows that, if 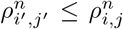 then 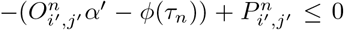 for all 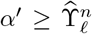. This implies that 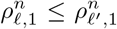 for all ℓ’ > ℓ. In plain English, each time that a α-bound 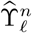 of Type 1 appears, there is a *x*-bound which is no longer required for defining 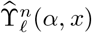 for all 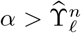.

The situation is symmetrical if 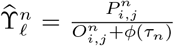. Each time that a α-bound 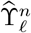 of Type 2 appears, there is a *x*-bound of *h_n_* which is no longer required for defining 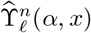 for all 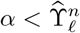.

Since all *x*-bounds used in the definition of *h_n_* belong to 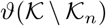 and can be involved at most once in the appearance of α-bounds of Type 1 and at most once in the appearance of α-bounds of Type 2, we get that the number of “new” α-bounds of 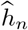 is smaller than 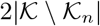.

### Corollary 1.

Under the notations of Theorem 2 and for all nodes n of 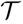, both *w_n_* and 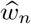 are smaller than 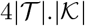

*Proof*. From Equation 4 and Theorem 3 from [5], if *n* is a child of the root then 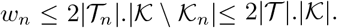. Let *m* is a non-root node of 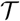, there exists a path *k*_1_, *k*_2_, … *k_ℓ_* of 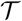 with 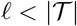 going from a child of the root ko to *k_ℓ_* = *m*. The number *w_m_* of *α*-bounds of *h_m_* is obtained by adding all the increases along the path *k*_1_, *k*_2_, … *k_ℓ_* to *w*_*k*_1__:

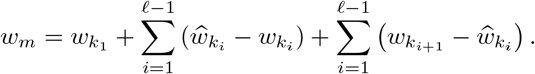

From Equation 4 and Theorem 3 from [5] and since *k_1_* is a child of the root, we have

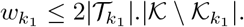

Proposition 1 ensures that

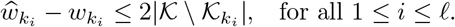

Equation 10 and Theorem 3 of [5] imply that

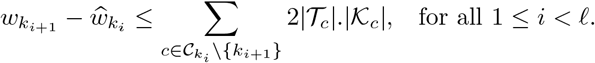

We get that

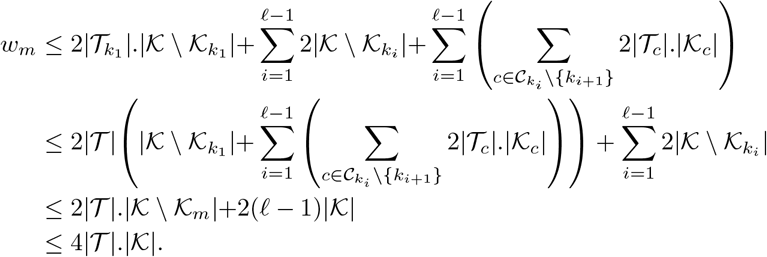

Since 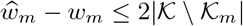 (Proposition 1), we have that

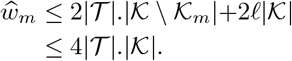

### Proposition 2.

*Under the assumption that the number of children of any node is bounded independently of the size of the tree, the algorithmic complexity of the computation of the cost functions h_n_ and* 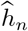 *for all nodes n of a tree* 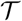 *with a set of known nodes* 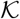 is 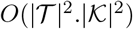 *both in time and memory space*.

*Proof*. From Theorem 1, less than 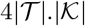 α-bounds are required for defining 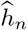. Over intervals between two successive α-bounds, 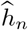 requires at most 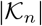 x-bounds. It follows that the total number of bounds required for defining 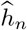 is 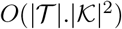. Since the part of Algorithm 1 computing 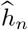 from 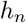 is linear with the total number of bounds required to define 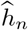, its complexity is 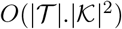 both in time and memory space.

The cost function *h_n_* is obtained by summing the partial cost functions 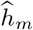 of its direct ancestor *m* and the partial cost functions 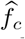 of its siblings c, which is performed by using a procedure similar to that of merging sorted lists. Under the assumption that the number of children *m* is bounded independently of the size of the tree, this operation is linear with the sum of all the bounds of these cost functions cost functions which is 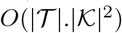.

In conclusion, the computation of all the partial cost functions of 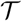 is 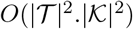 both in time and memory space.

## 4 Integrating parsimonious costs over asymmetry parameter(s)

### 4.1 Single parameter case: no-split cost

We show here how to sum the parsimonious *α*-cost 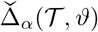 over all asymmetry parameters *α*. Let us start by recalling Remark 3 from [5].

#### Remark 1 ([5]).

Let *r* be the root of 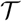 and 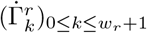 be the elements of

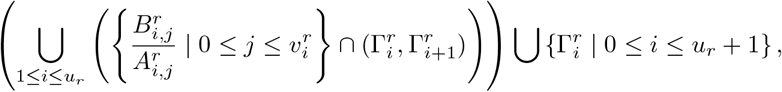

indexed in increasing order. There exists a sequence 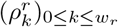 of increasing values in 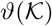 such that for all *0 ≤ k ≤ w_r_* and all 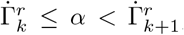 α-parsimonious reconstruction associates 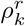 to *r*. If 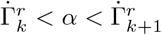 then all α-parsimonious reconstructions associate 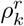 to *r*.

An algorithm computing 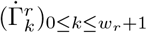 and 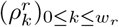 from *f_r_* is provided in [5]. Its complexity is linear with the total number of bounds required for defining *w_r_*.

From Theorem 1, we have that the most parsimonious *α*-cost is, for all 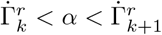,

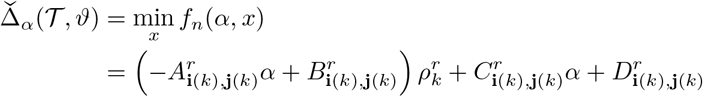

where ***i**(k)* is the index *i* such that 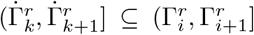 and ***j**(k)* is the index *j* such that 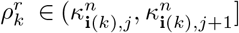.

Integrating the parsimonious cost of 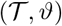 over all possible values of the asymmetry parameter can then be done explicitly:

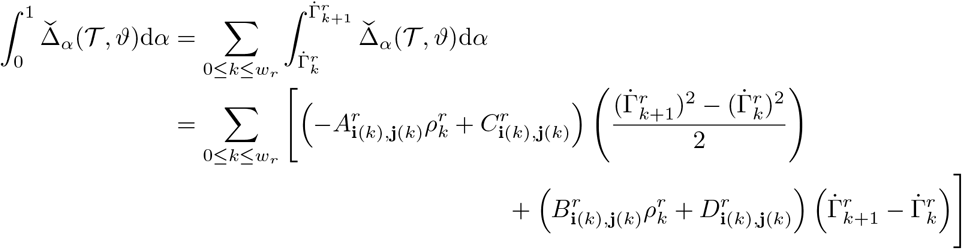

The quantity such obtained will be referred to as the *no-split cost* of 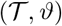.

### 4.2 Two parameters case: A - and B-split costs

In order to test a trend change at a node *n* of 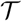, we shall consider the two ways of splitting 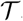 at *n* displayed in Figure 1, namely, the most parsimonious costs obtained with

(A) an asymmetry parameter *α’* on the subtree 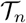 and an asymmetry parameter *α* on the rest of 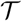 (i.e., with *α’* on the red part and with *α* on the blue part of Figure 1-left).
(B) an asymmetry parameter *α’* on the subtree consisting of the direct ancestor of n the branch bearing *n* and 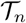 (i.e., the stem subtree of n) and an asymmetry parameter α on the rest of 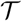 (i.e., with *α’* on the red part and with α on the blue part of Figure 1-right);

In Case A, given two asymmetry parameters *α* and *α’*, we are able to compute the smallest cost which can be achieved by associating *x* to *n* with asymmetry parameters α’ on 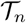 and *α* on the rest of 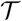. Namely, this cost is

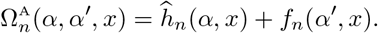

In the same way for Case B, the smallest cost which can be achieved by associating *x* to the direct ancestor of *n* and by using the asymmetry parameter *α’* on the stem subtree of *n* and *α* on the rest of 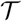 is given by

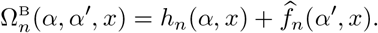

We shall detail only Case A, Case B being very similar.

From Theorems 1 and 2, the map 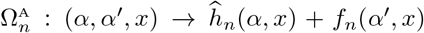 is piecewise linear and continuous. Namely, under the notations of these theorems, for all 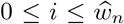, all *0 ≤ j ≤ u_n_* and by putting 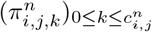. for the elements of 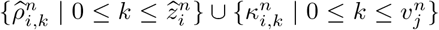 sorted in increasing order, we have that, for all 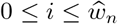, all 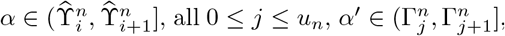, all 0 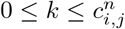 and all 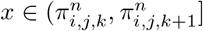,

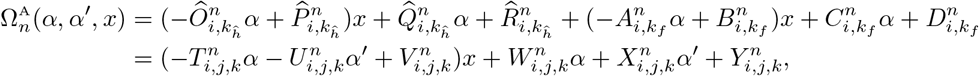

where 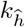 is the index such that 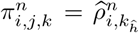 is the index such that 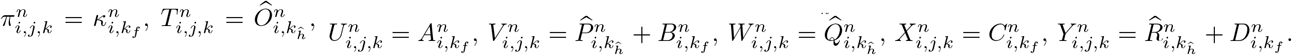.

Let us remark that, from Theorems 1 and 2, for all asymmetry parameters *α* and *α’*, the map 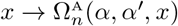 is convex.

The smallest cost obtained with asymmetry parameter *α’* on 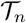 and with asymmetry parameter *α* elsewhere is

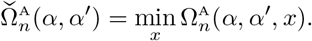

Since 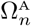 is convex with respect to *x*, for all 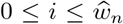, all 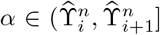, all 0 ≤ j ≤ u_n_ and all 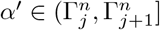, we have that

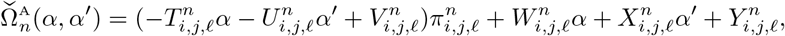

where *ℓ* is the smallest index such that 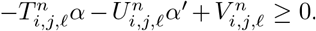. Such an index *ℓ* always exists since Theorems 1 and 2 imply that 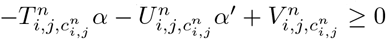.

For all indexes 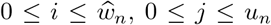, the lines of equation 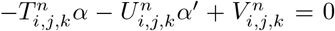 with 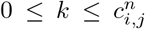 partition 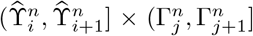 into subsets on which the smallest index *ℓ* such that 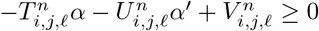 remains the same, i.e. in which 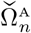 is linear with respect to *α* and *α’* (see Figure 2 without taking into account the dotted line).

**Figure 2:**
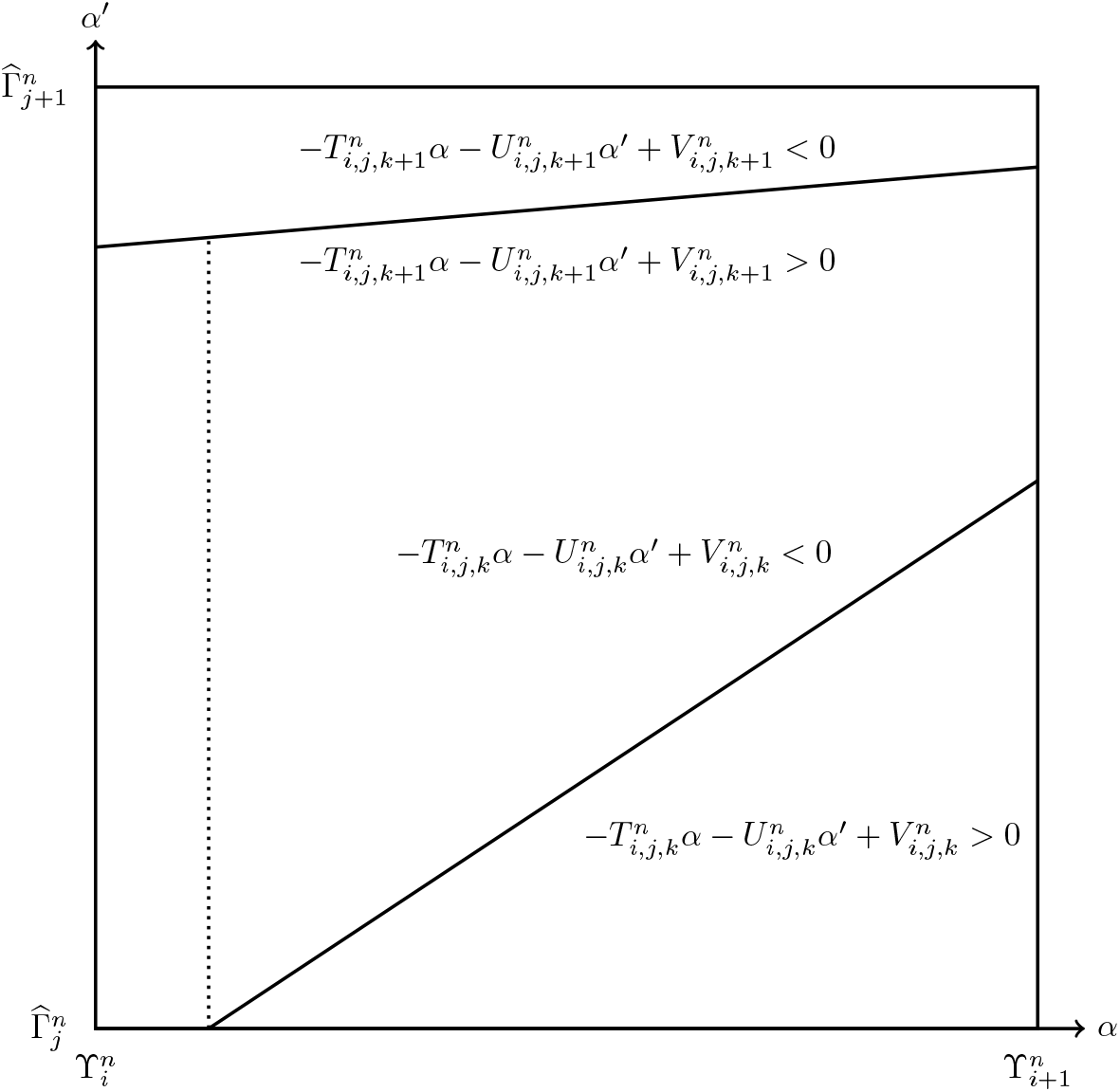
Partition of 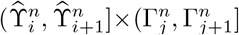 resulting from the lines of equations 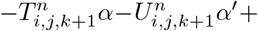 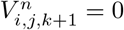 and 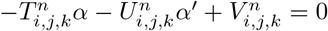 (the top and the bottom solid lines respectively).

#### Proposition 3.

*For all indexes* 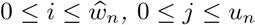, *if there exists* 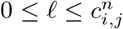 *such that* 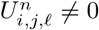 *and such that the line of equation* 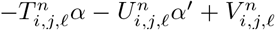 *intersects* 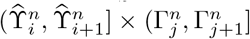 *then the points (α, α’) for which* 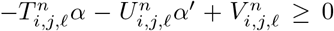 *are those which are below the line of equation* 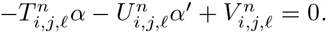.

*Proof*. Since from Theorem 1, 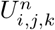 is nonnegative for all 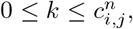, we have that 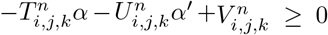 if and only if 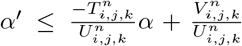, i.e., the point *(α, α’)* is below the line of equation 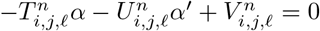 (Figure 2).

#### Proposition 4.

*For all indexes* 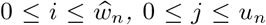 and 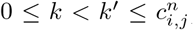, *the lines of equations* 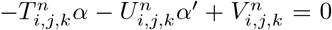 *and* 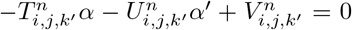 *do not intersect each other at any point in* 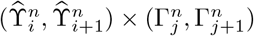.

*Proof*. Since 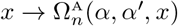 is convex, we have that 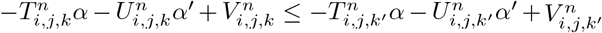. It follows that 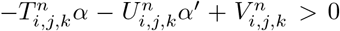 implies 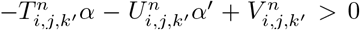 for all 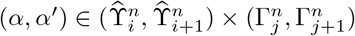.

From the implicit assumption that all the bounds of partial functions are required, if *k ≠ k^’^* then 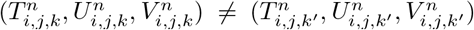, i.e., two lines intersect each other at a single point. If the; lines of equations 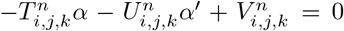 and 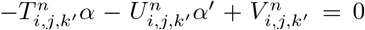 intersect each other at a point of 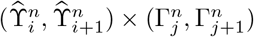, they split 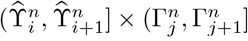 into four non-empty parts among which at least one contains points (*α, α’*) such that 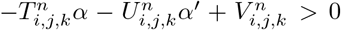 and 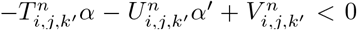 which is in contradiction with the convexity of 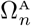 with respect to *x* since we assume *k* < *k*’.

#### Proposition 5.

Let *i* and *j* be two indexes with 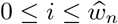 and 0 < *j* < *u_n_*.

a. If there exists 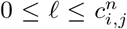. such that both 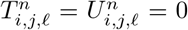 and 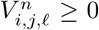 then there is no line of equation 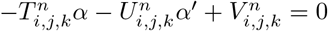 with *k > ℓ* intersecting 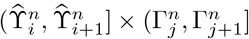.
b. If there exists 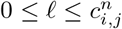. such that 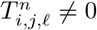 and 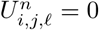 then

- if 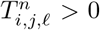 and 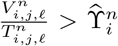 then there is no line of equation 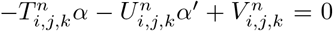 *with k> ℓ intersecting* 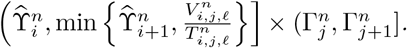.
- if 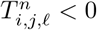 and 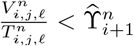 then there is no line of equation 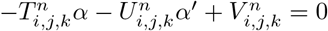 with *k >ℓ* intersecting 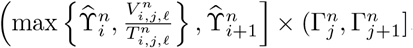.
c. For all 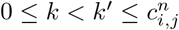, the part of the line corresponding to *k’* intersecting 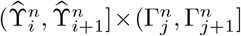 is above that of *k* on 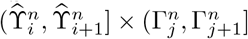.

*Proof*. These three properties are again consequences of the convexity of the map 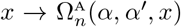 for all 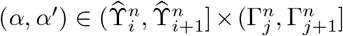, i.e., of the fact that the sequence 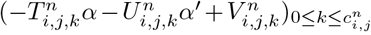. increases with *k*. In particular, this implies that if there exist a part 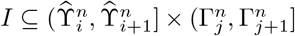 and an index *ℓ* is such that 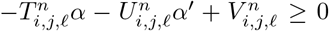 for all (*α, α’*) *∈ I* then there is no line of equation 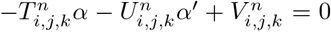 with *k > ℓ* splitting *I* into two non-empty parts, since this would imply the existence of points *(α, α’) ∈ I* with 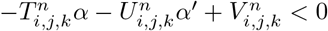.

The three assertions are proved as follows.

a. If 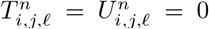 and 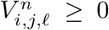 then we have that 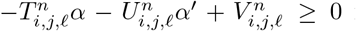 for all 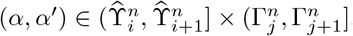.
b. Let us assume that 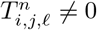 and 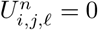

- If 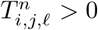 0 and 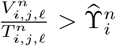 then we have that 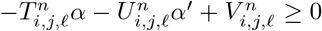 for all 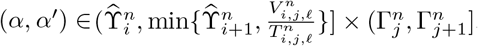.
- If 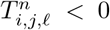 and 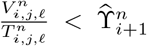 then we have that 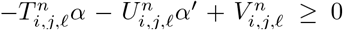 for all 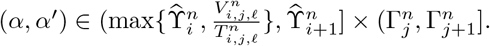
c. The property directly follows from Proposition 3 and from the opening remark of this proof.

The *A-split cost of* 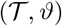 *at n* is the integral of 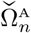 over all pairs *(α, α’) ∈ [0,1]*^2^:

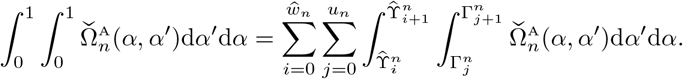

For all 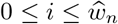 and all *0 < j < u_n_*, computing

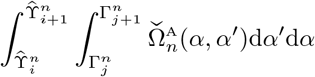

can be performed by splitting 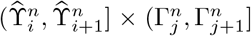 into trapezoids of the form {(α, α’) | *α* < *α* < *w* and ca + b < a’ < *ca* + *d}* (i.e., with two vertical sides) in which the coefficients of 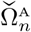 are constant and of the form:

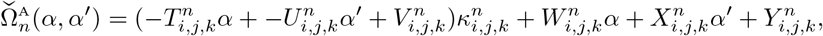

for a certain index *k*. The different lines of Figure 2, here including the dotted one, partition 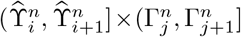 into such trapezoids (the triangle below the line of equation 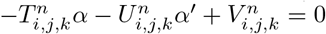 is seen as a degenerated trapezoid with a left side of length 0).

The algorithm partitioning 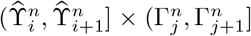 into trapezoids is not presented here since it is quite basic thanks to Propositions 3, 4 and 5 which limit the number of situations that we have to deal with.

The integral of 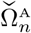 over a trapezoid *{(α, α’) | β < α < w* and *αα + b < α’ < cα + d}* can be explicitly computed:

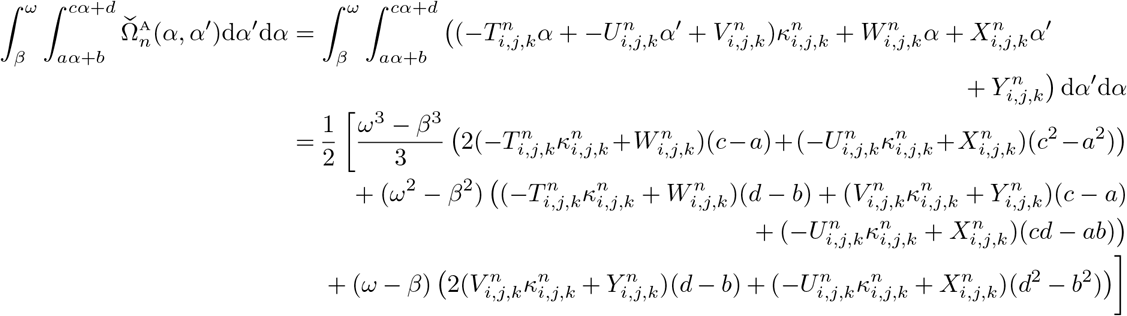

Symmetrically, by setting

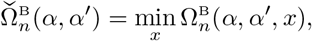

We define the *B-split cost of* 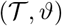 at n as

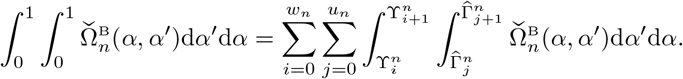

All the remarks stated above about A-split costs still hold for B-split costs which are computed in the very same way.

## 5 Applications

For all examples and datasets below, the parsimonious splits identification was performed by setting 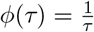 for computing the evolutionary costs.

### 5.1 Two small examples

Figure 3 and Table 1 illustrate the fact that considering more than one asymmetry parameter on a tree may or may not lead to lower the parsimonious cost. The two examples displayed in Figure 3 are based on the same tree but with two different initial functions (in brackets just after idents of the tips). Table 1 displays the A - and B-split costs for all nodes of the tree with regard to the character values of Figure 3.

**Figure 3:**
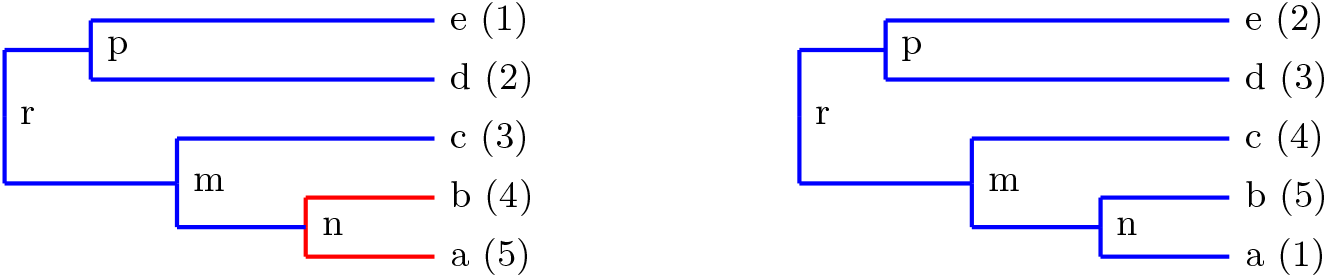
A tree with two sets of character values (in brackets) for tips. Left: A A-split at node ‘n’ improves the parsimonious cost of the whole tree. Right: no split improve the parsimonious cost of the whole tree (see Table 1).

**Table 1:**
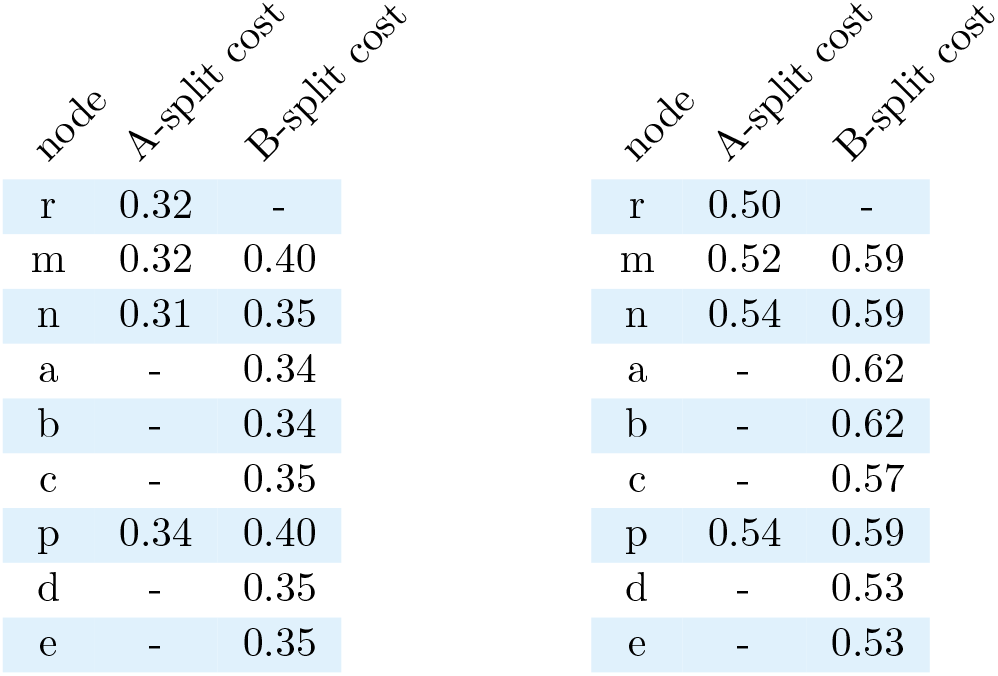
Left: (resp. Right:) A - and B-split costs for all nodes of the tree with character values of Figure 3-left (resp. of Figure 3-right). The first line of the tables displays the A-split cost of the root ‘r’, i.e., the no-split cost computed as presented in Section 4.1.

| node | A-split cost | B-split cost |
| --- | --- | --- |
| r | 0.32 | - |
| m | 0.32 | 0.40 |
| n | 0.31 | 0.35 |
| a | - | 0.34 |
| b | - | 0.34 |
| c | - | 0.35 |
| p | 0.34 | 0.40 |
| d | - | 0.35 |
| e | - | 0.35 |
| r | 0.50 | - |
| m | 0.52 | 0.59 |
| n | 0.54 | 0.59 |
| a | - | 0.62 |
| b | - | 0.62 |
| c | - | 0.57 |
| p | 0.54 | 0.59 |
| d | - | 0.53 |
| e | - | 0.53 |

In the example at the left of the figure, the A-split cost at node ‘n’ is smaller than the no-split cost (i.e., the A-split at the root ‘r’ - first line of Table 1-left). Conversely, in the example at the right, there is no A - or B-split cost lower than the no-split cost.

From a general point of view, it may be worth paying interest not only on the most parsimonious split but also on the other splits with costs smaller than the no-split cost.

### 5.2 Two biological datasets

Figures 4 and 5 display all the splits that lead to a cost lower than the no-split one. Splits are represented with ‘× ‘ at the corresponding nodes. Figures also display the ranks of the splits with regard to their costs (ordered from lowest to highest). The color of the subtree pending from a split depends on the corresponding split cost, more precisely on the cost of the “most recent” split including it. The color scale starts from blue which corresponds to the no-split cost or higher to red for the lowest cost observed. These two figures were output by our software *ParSplit*.

**Figure 4:**
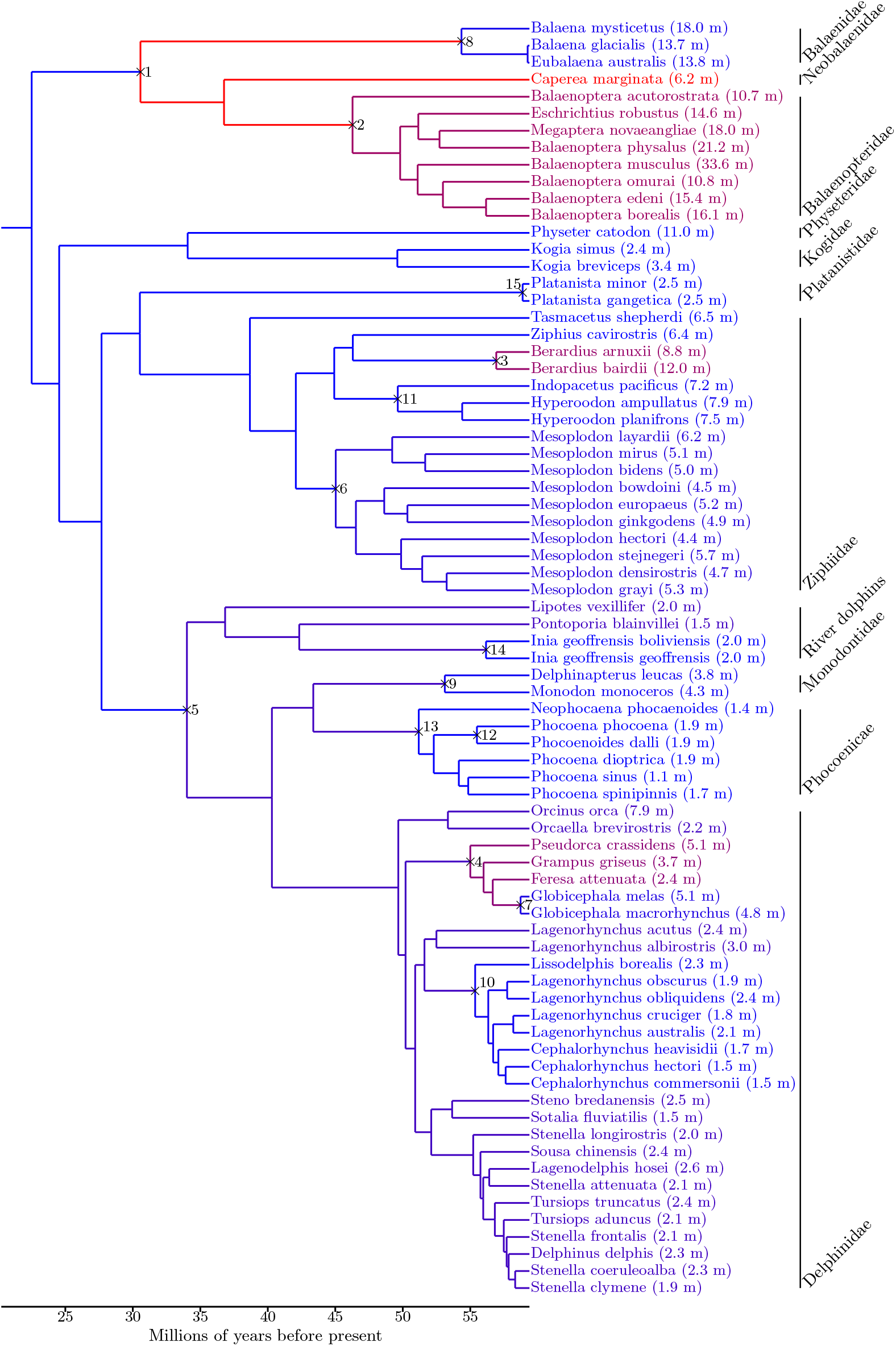
Parsimonious splits of the tree of cetaceans according to their body sizes with their ranks. All the parsimonious splits are A-splits. Dataset from [18]

**Figure 5:**
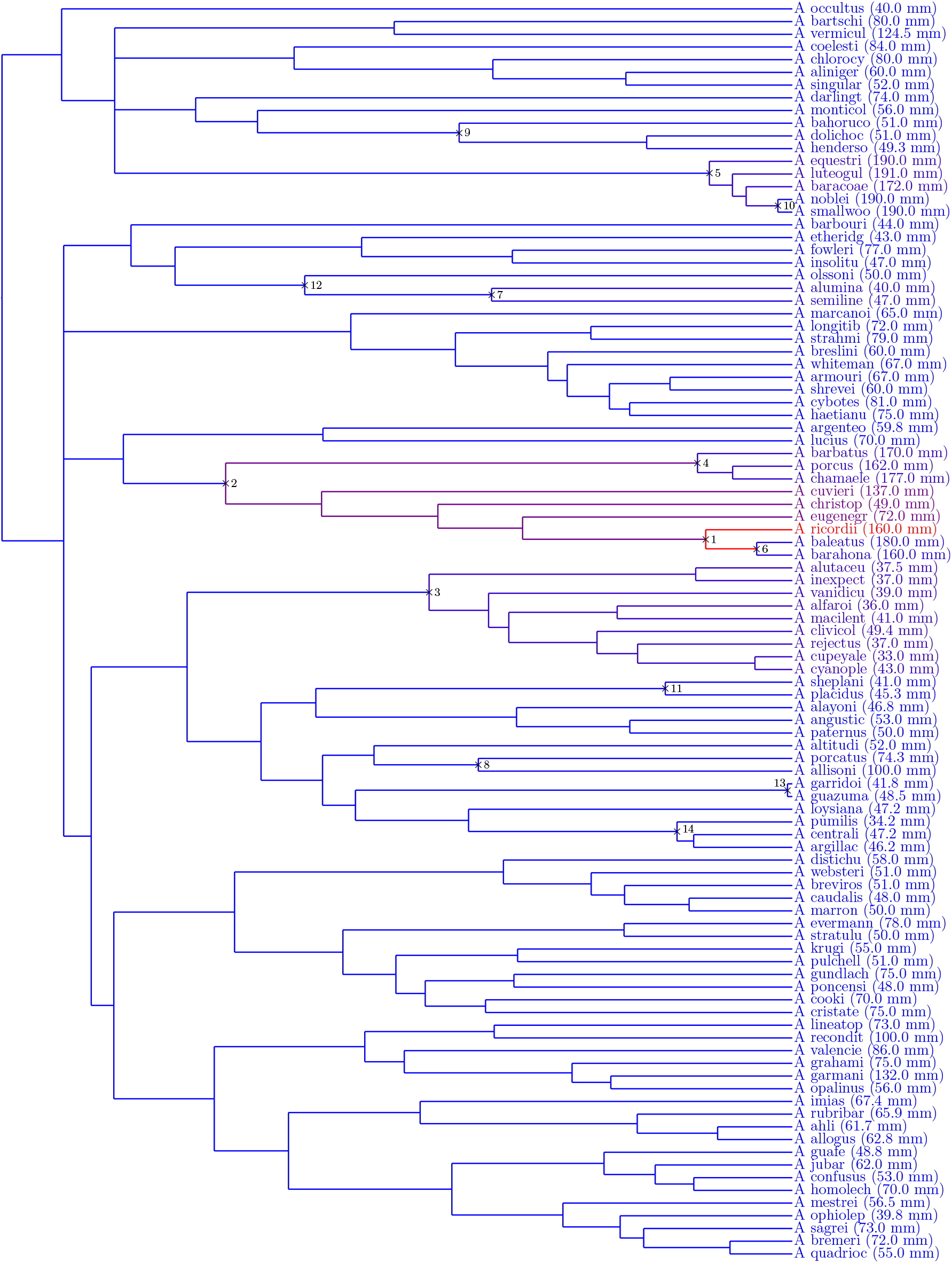
Parsimonious splits of the tree of *Anolis* lizards according to their body sizes with their ranks. All the parsimonious splits are A-splits. Dataset from [20]

For the two datasets studied here, all the most parsimonious splits are A-splits. Moreover, no B-split costs were smaller than the no-split cost. This is not a general behavior since we did encounter cases in which some of the B-split costs were lower than than the no-split cost and even smaller than the A-split costs at the same nodes.

We observe in Figure 4 that the split with the lowest cost occurs at the most recent common ancestor of the mysticetes (i.e., *Balaenidae, Neobalaenidae* and *Balaenopteridae*). First, this split is consistent with a probable shift in diet type, since *mysticetes* are filter feeders. Second, authors of [18] also distinguished this clade and found that “The largest phylogenetic mean body size was recovered for *Mysticeti*…” by using approaches based on heterogeneous stochastic models from [21, 22].

Figure 5 displays the parsimonious splits of the tree of *Anolis* lizards according to the body size of males. This dataset comes from [20] in which authors used two methods for identifying rate shifts both based on stochastic models. With the first one, they detected two shifts at the same nodes as those of our two splits with the smallest costs, ranked in the same order as with our approach. With their second approach, they identified a third shift at the same node as that of the split ranked 5 with our method for which the costs of split of ranks 3, 4 and 5 are very close.

### 5.3 Discussion

It is worth noting that, as observed on the two biological datasets above, a parsimony-based approach may identify the evolutionary shifts detected from methods based on stochastic models of evolution. In all the situations where parsimony is preferred, our approach provides a parsimonious solution for identifying evolutionary shifts.

We emphasize the fact that the different splits displayed in Figures 4 and 5 have to be considered as different alternatives for splitting the tree into two parts and absolutely not as a split of the tree into more than two parts. Finding the most parsimonious split of the tree into k > 2 parts in the same sense as above could be performed with ideas similar as those developed in this work. The algorithmic complexity increases exponentially with k but computations would be still feasible for small k (e.g. smaller than 5 for usual phylogenetic trees). In particular, one could seek for the most parsimonious cost obtained with a same asymmetry parameter on k subtrees and another parameter for the rest of the tree, following the ideas of [10] for studying convergence with Ornstein-Uhlenbeck processes.

